# The hyperstability and composition of *Giardia*’s ventral disc highlights the remarkable versatility of microtubule organelles

**DOI:** 10.1101/361105

**Authors:** C. Nosala, K.D. Hagen, T.M. Chase, K. Jones, R. Loudermilk, K. Nguyen, S.C. Dawson

## Abstract

*Giardia* is a common protistan parasite that causes diarrheal disease worldwide. Motile trophozoites colonize the small intestine, attaching to the villi with the ventral disc, a unique and complex microtubule (MT) organelle. Attachment to the host epithelium allows *Giardia* to resist peristalsis during infection of the host gastrointestinal tract. Despite our emerging view of the complexity of ventral disc architecture, we are still in the very preliminary stages of understanding how specific structural elements contribute to disc stability or generate forces for attachment. The ventral disc is a large, dome-shaped, spiral MT array decorated with microribbon-crossbridge protein complexes (MR-CB) that extend upward into the cytoplasm. To find additional disc-associated proteins (DAPs), we used a modified method for disc biochemical fractionation in high salt followed by shotgun proteomic analyses and validation by GFP-tagging. Using this method in conjunction with an ongoing subcellular localization screen, we identified 54 new DAPs. Of the 87 DAPs confirmed to date, 54 localize only to the disc, and the remainder localize to additional structures including the flagella, basal bodies, or median body. Almost one third of the known DAPs lack any homology to proteins in other eukaryotes and another one third simply contain ankyrin repeat domains. Many DAPs localize to specific structural regions of the disc, including the ventral groove region and disc margin. Lastly, we show that spiral singlet MT array comprising the disc is hyperstable and lacks dynamic instability, and we attribute these unique properties to the presence of both novel DAPs as well conserved MAPs and MIPs that are known to stabilize ciliary doublet and triplet MTs.

## Introdcution

Many microbial eukaryotes possess complex interphase microtubule organelles, whose structures offer a unique perspective into how the seemingly boundless capabilities of microtubule polymers are used to generate diverse forms and functions in cells. As compared to the more well-studied MT-based mitotic spindle or cilium (Chaaban and Brouhard, 2017), microtubules in these unique organelles are often arranged into elaborate and stable higher-order assemblies and are composed of proteins that lack homology to MT-binding proteins in other eukaryotes (Hagen et al., 2011; Hu et al., 2006; Preisner et al., 2016). Interphase microtubule-based organelles in protists thus represent an untapped reservoir of non-canonical MT-binding proteins governing MT assembly, nucleation, or dynamics (Dawson and Paredez, 2013).

The widespread protistan intestinal parasite *Giardia lamblia* is defined by one such elaborate cup-shaped microtubule (MT) organelle, the ventral disc (Crossley and Holberton, 1983a; Crossley and Holberton, 1985; Feely et al., 1982; Friend, 1966; Holberton, 1973; Holberton, 1981). Infectious *Giardia* cysts are commonly ingested from contaminated water sources (Einarsson et al., 2016) and excyst into flagellated trophozoites. Using the ventral disc, trophozoites attach to the intestinal microvilli and resist being dislodged by peristalsis (Elmendorf et al., 2003; Nosala and Dawson, 2015). Trophozoite colonization leads to acute or chronic diarrheal disease in humans and other animals (Nosala and Dawson, 2015). While the molecular mechanisms of attachment are not well understood, *Giardia* trophozoites are thought to attach to surfaces via hydrodynamic suction-based forces that result in a pressure differential underneath the disc relative to the outside medium (Hansen et al., 2006; Holberton, 1974). Overall conformational changes of regions of the disc may also either create or modulate attachment forces in the absence of canonical MT dynamic instability (Dawson et al., 2007; Woessner and Dawson, 2012).

The intricate architecture of the ventral disc was first described by Cheissin over 50 years ago (Cheissin, 1964). Early work described the novel structural elements associated with the ventral disc microtubules (Friend, 1966), and the first 3D high-resolution structure of the ventral disc was obtained recently using cryo-electron tomography (cryo-ET) (Schwartz et al., 2012). Cryo-ET of whole isolated ventral discs with sub-tomographic averaging defined an elaborate cytoskeletal architecture with dense protein complexes coating nearly every protofilament of the microtubule spiral array (Figure 1A and (Schwartz et al., 2012)).

**Figure 1.**
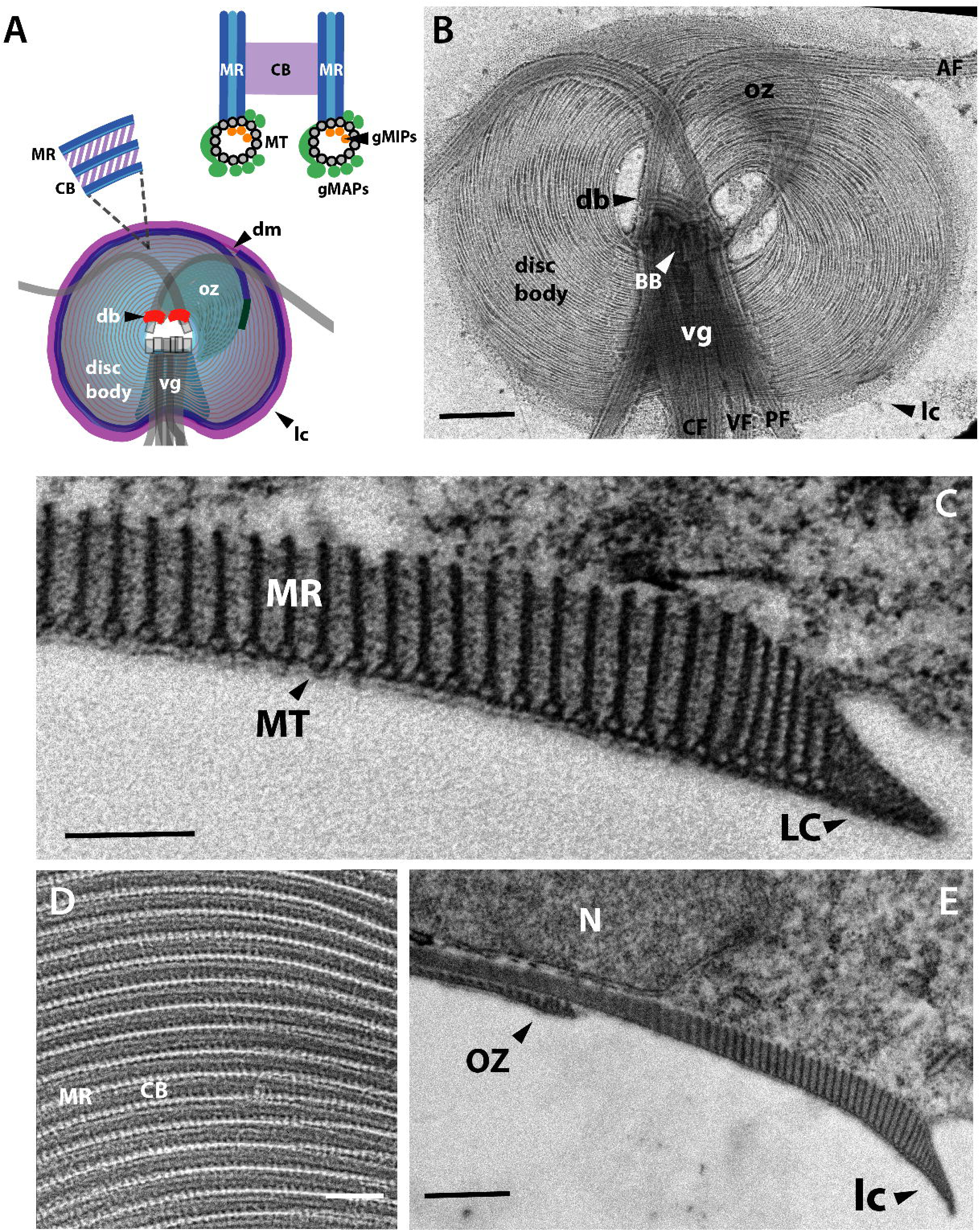
Elaborate and unique protein complexes are associated with the disc spiral singlet microtubule array. A schematic of the ventral disc indicating the primary structural elements is shown in panel A: disc body, oz: overlap zone, lc: lateral crest, vg: ventral groove region, db: dense bands, dm: disc margin (A), MT: microtubule, MR: microribbon, CB: crossbridge (B,C). The inset shows a cross section of the MR/CB complex and the Giardia MAPs (gMAPs, green) or MIPs (gMIPs, orange). In B, a negative-stained cytoskeletal preparation of the ventral disc highlights the primary structural elements as well as the eight cytoplasmic axonemes (CF = caudal flagella; VF = ventral flagella; PF = posteriolateral flagella; AF = anterior flagella) and basal bodies (BB). Scale = 1 µm. In C, transmission electron microscopy of thin sections of whole embedded trophozoites show both microribbons (MR) associated with each MT of the spiral array and the associated lateral crest (LC). Scale = 200 nm. Negative staining of the disc (D) highlights the microribbons (MR) linked with regularly spaced crossbridges (CB). Scale = 200 nm In E, the overlap zone (OZ) of the MT spiral array, along with the MR-CB complexes are shown in cross section (N = nucleus). Scale bars = 500 nm.

The complex architecture, composition, and functioning of the ventral disc exemplifies the profound capabilities of microtubule polymers. Roughly one hundred, uniformly spaced MTs spiral one and a quarter turns to form an 8 µm in diameter domed organelle required for host attachment (Brown et al., 2016). An overlap zone occurs between the upper and lower portions of the spiral (Figure 1A, B). Associated with the entire length of the MT spiral are unique repetitive substructural elements – the trilaminar microribbons – that extend 150-400 nm dorsally into the cytoplasm and are believed to stabilize the overall disc structure (Holberton, 1973; Holberton, 1981) (see Figure 1C, D). Regularly spaced “crossbridge” structures link adjacent microribbons (Holberton, 1973) and repeat every 16 nm (Figure 1D), corresponding to the distance of two alpha/beta tubulin dimers. MAPs termed sidearms and paddles repeat every 8 nm and are spaced at the distance of a single alpha/beta tubulin dimer (Figure 1). Other repetitive MT-associated substructures include three MT-associated proteins (gMAPs 1-3) and three MT inner proteins (gMIPs 5, 7 and 8) (Schwartz et al., 2012). An associated structure at the disc periphery, the lateral crest (Figure 1C), forms a seal with surfaces in early attachment (Feely, 1990; Feely et al., 1982; House et al., 2011).

Despite our emerging view of the complexity of the ventral disc architecture (Brown et al., 2016), we still know very little about the composition of the disc or the contribution of specific structural elements to disc stability and the generation of attachment forces. While abundant MT-associated proteins termed ‘giardins’ were isolated from the disc two decades after the initial disc structures were described (Crossley and Holberton, 1983a), the identities of proteins composing the unique MT-associated structure of the disc have remained elusive. In a comprehensive proteomic analysis of detergent-extracted, isolated ventral discs, we previously identified nearly twenty new disc-associated proteins (now referred to as DAPs, rather than giardins) that localize to regions of the ventral disc or lateral crest (Hagen et al., 2011).

The complexity of the disc architecture as revealed by more detailed cryo-ET implies that there are disc proteins still to be discovered (Brown et al., 2016). Here we have identified 54 additional disc-associated proteins (DAPs) using a modified method of disc biochemical fractionation in high salt followed by shotgun proteomic analyses with subsequent confirmation of these or other DAPs by GFP-tagging (Harb and Roos, 2015). Fifty-four of the 87 total DAPs localize exclusively to the disc, and the remainder localize to other MT structures such as the flagella, basal bodies, or median body. Lastly, we demonstrate that the disc singlet MT spiral array is a “hyperstable” structure that includes conserved MAPs and MIPs (Nosala et al., 2018) that are known to stabilize ciliary double and triple MTs (Ichikawa and Bui, 2018; Stoddard et al., 2018). Many of the newly described DAPs likely facilitate disc MT nucleation, MT plus-end binding, or stabilization of the curved spiral array of microtubules. Future genetic and functional analyses of DAPs will be central toward understanding disc architecture, assembly and dynamics.

## Materials and Methods

### *Giardia* culture conditions

*Giardia* strains were cultured in sterile 16 ml screw-capped disposable tubes (BD Falcon) containing 12 ml modified TYI-S-33 medium supplemented with bovine bile and 5% adult and 5% fetal bovine serum (House et al., 2011). Cultures were incubated upright at 37°C without shaking and reached confluence at ~48 hours. Puromycin (50 µg/ml) was added when required to maintain episomal plasmids in GFP strains.

### C-terminal GFP tagging of candidate disc-associated proteins

*Giardia* GFP strains were created using Gateway cloning as previously described (Hagen et al., 2011). To add C-terminal GFP tags to candidate DAPs, PCR forward primers were designed to bind 200 bp upstream of the gene to include the *Giardia* native promoter. Full-length genes lacking stop codons were amplified from *Giardia lamblia* strain WBC6 genomic DNA using either PfuTurbo Hotstart PCR Master Mix or Easy-A High Fidelity PCR Master Mix (Agilent) and were cloned into the ThermoFisher Scientific Gateway entry vectors pENTR/D-TOPO (blunt-end, directional TOPO cloning) or pCR8/GW/TOPO (for efficient TOPO TA cloning), respectively. Inserts were sequenced to confirm gene identity and correct orientation. To generate DAP-GFP fusions, entry vectors were recombined with the *E. coli*/*Giardia* Gateway destination vector pcGFP1F.pac (GenBank #MH048881) (Dawson and House, 2010) via LR recombination using LR Clonase II Plus (ThermoFisher Scientific). Clones were screened by AscI digest and Qiagen’s Plasmid Plus Midi Kit was used to prepare bulk plasmid DNA. To create *Giardia* strains expressing GFP-tagged candidate DAPs, 20 µg plasmid DNA was introduced into 10^7^ *G. lamblia* strain WBC6 (ATCC 50803) trophozoites by electroporation using a Bio-Rad GenePulserXL as previously described (Dawson et al., 2007). Transfectants were initially selected with 10 µg/ml puromycin; the antibiotic concentration was increased to 50 µg/ml once cultures reached 50% confluence (typically 7 to 10 days). Cultures were maintained with selection for at least two weeks prior to the preparation of frozen stocks.

### Biochemical fractionation of the Giardia cytoskeleton

Detergent extraction of *Giardia’s* microtubule cytoskeleton was performed as previously described (Hagen et al., 2011) with modifications indicated below. Medium was decanted from each confluent culture tube and trophozoites were washed *in situ* three times with 5 ml of warmed 1X HEPES Buffered Saline (HBS). To demembranate cytoskeletons, cells were iced for 15 minutes in the last 1X HBS wash, pelleted by centrifugation (1000 × g, 5 minutes), and resuspended in 1 ml of 1X PHEM (60 mM PIPES, 25 mM HEPES, 10 mM EGTA, 1 mM MgCl2, pH 7.4) containing 1% Triton X-100 and 1M KCl. 1X HALT protease inhibitor cocktail (Roche) was added to prevent proteolysis. This solution was transferred to an Eppendorf tube and vortexed continuously for 30 minutes at medium speed. Ventral disc cytoskeletons were then pelleted as above and washed twice in 1X PHEM lacking both Triton X-100 and KCl. Sufficient extraction of cytoskeletons was confirmed by wet mount using DIC microscopy.

Cytoskeletons were fractionated as previously described (Crossley and Holberton, 1983a; Crossley and Holberton, 1983b; Crossley and Holberton, 1985) and Figure 2A). Following detergent extraction, an aliquot of demembranated cytoskeletons in PHEM was retained as ‘P1.’ Cytoskeletons were then pelleted as described above, washed, and resuspended in CB buffer (10mM Tris, 1mM EDTA, pH 7.7) for 48 hours to further disrupt the cytoskeleton. Leftover cytoskeletal complexes were pelleted at 1000 x g for 5 minutes and the supernatant was retained as ‘S2.’ Complexes were next washed and resuspended in MR buffer (10mM HEPES, 5mM EDTA, pH 8.7) for 48 hours. Any remaining complexes were then pelleted, and the supernatant was retained as ‘S3.’ Complexes in the final pellet were resuspended in 1X PHEM and retained as ‘P3.’

**Figure 2:**
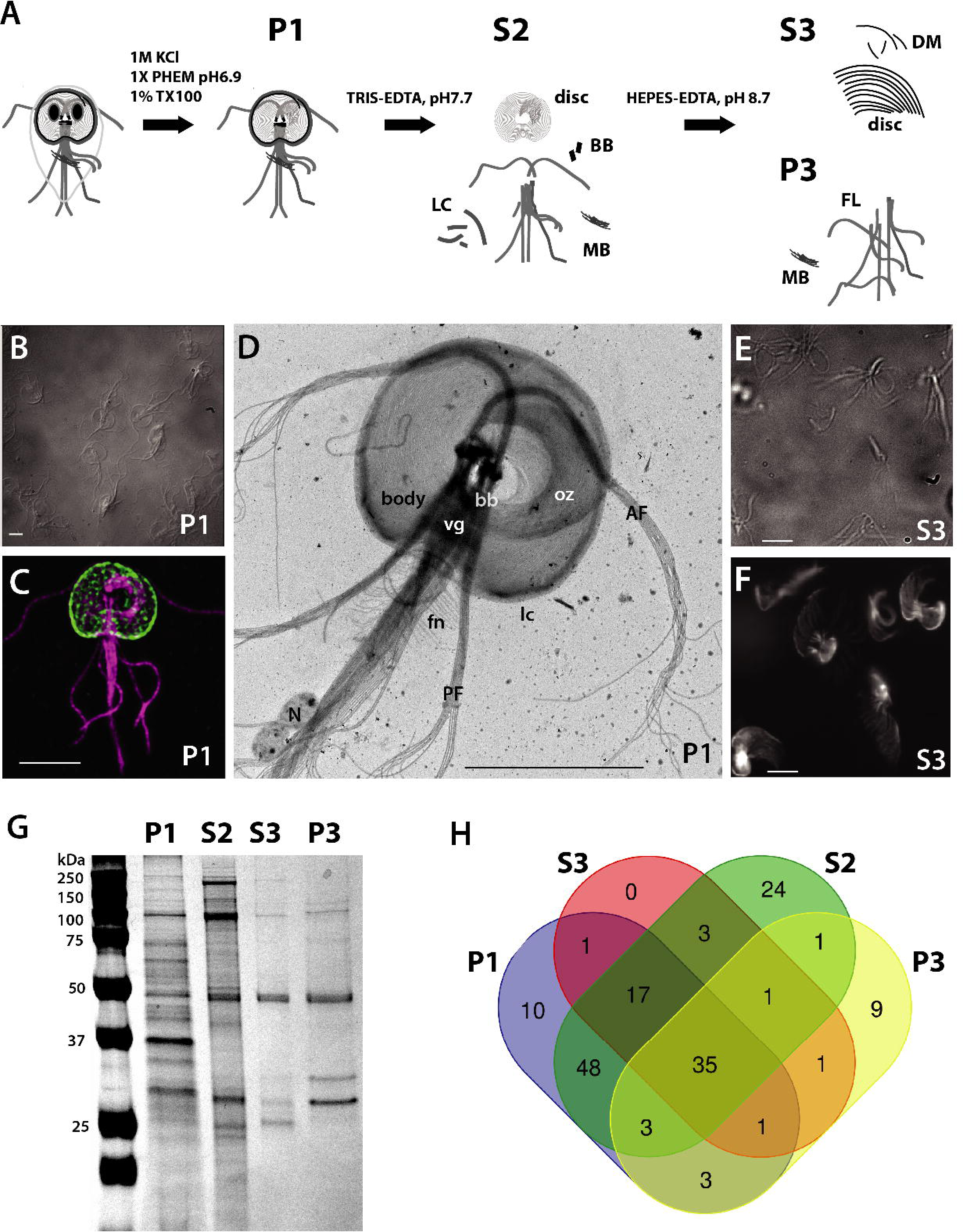
Biochemical fractionation readily extracts disc and axoneme proteins for shotgun proteomic analysis and mass spectrometry. Extraction of Giardia trophozoites with detergent and high salt (A) removed membrane and cytosol from the microtubule cytoskeleton (P1). Subsequent extractions with Tris-EDTA and HEPES further disrupted the disc resulting in the S2, P2, S3 and P3 fractions. Immunostaining of P1 fraction (B, C) shows that the disc (green, anti-delta-giardin) and flagella (purple, anti-alpha-tubulin) were retained. Scale = 1 µm (B). Scale = 5 µm (C). Negative staining of the P1 fraction (D) confirms the loss of the nuclei (N) and retention of the disc (body, vg, oz, lc), flagella (AF, PF) and the funis (fn). Scale = 5 µm. Greater dissociation of the disc from the axonemes is evident in the S3 fraction as shown by delta-giardin immunostaining (E, F). SDS-PAGE resolves proteins that are enriched in each pellet (P1, P3) and supernatant (S2, S3) fraction. Following mass spectrometry of each fraction (P1, P3 and S2, S3), the Venn Diagram comparison indicates some overlap between proteins in the various fractions (H). Scale bars = 5 μm.

### SDS-PAGE and Western Blotting

For Western blots, total cell lysate in RIPA buffer (20 μg) was run on Precise Tris·HEPES SDS PAGE gels, 4–20% (ThermoFisher). Proteins were transferred onto 0.45 micron nitrocellulose membrane (Bio-Rad) and blocked for 1 hour at room temperature in 5% (wt/vol) milk in PBS with 0.05% Tween. Antibodies were diluted in 5% (wt/vol) milk in PBS with 0.05% Tween as follows: anti-HA (Sigma) 1:5000, goat anti-mouse IgG HRP (Bio-Rad) 1:10000. Amersham ECL Prime substrate (GE Healthcare) was used for chemiluminescent detection. For SDS-PAGE of fractionated proteins, 20 ul of total undiluted fraction samples were run as for Western blots. Gels were stained in True Blue for 1 hour, then rinsed with MilliQ water and incubated with MilliQ water overnight at room temperature to destain.

### Proteomic analyses of fractions and mass spectrometry

All MS/MS samples were analyzed using X! Tandem (The GPM, thegpm.org; version X! Tandem Alanine (2017.2.1.4)). X! Tandem was used to search the uniprotgiardiaintestinalis_Craprev database (14528 entries) assuming the digestion enzyme trypsin and with a fragment ion mass tolerance of 20 PPM and a parent ion tolerance of 20 PPM. X! Tandem variable modifications specified were Glu->pyro-Glu of the N-terminus, ammonia-loss of the N-terminus, gln->pyro-Glu of the N-terminus, deamidated of asparagine and glutamine, oxidation of methionine and tryptophan and dioxidation of methionine and tryptophan. Scaffold (version Scaffold_4.8.4, Proteome Software Inc., Portland, OR) was used to validate MS/MS based peptide and protein identifications. Peptide identifications were accepted if they exceeded specific database search engine thresholds. Protein identifications were accepted if they contained at least 5 identified peptides. Proteins that contained similar peptides and could not be differentiated based on MS/MS analysis alone were grouped to satisfy the principles of parsimony. Proteins sharing significant peptide evidence were grouped into clusters.

### Live imaging of GFP-tagged DAP strains

Strains were thawed from frozen stocks and cultured for 24 to 48 hours prior to live imaging. Trophozoites were iced 15 minutes, harvested by centrifugation at 1000 x g, resuspended in warm medium, and incubated in 96-well black glass bottom imaging plates (Cellvis, Mountain View, CA) for up to 2 hours at 37°C in a nitrogen-enriched atmosphere to promote attachment. The medium was replaced with 1X HBS, and the trophozoites were incubated under the same conditions for 30 minutes. Additional warm HBS washes were performed as needed during imaging to remove detached cells. Imaging was performed using differential interference contrast (DIC) and epifluorescence with a Leica DMI6000B wide-field inverted fluorescence microscope (Plan Apo 100X, NA 1.40 oil immersion objective). Optical sections were acquired at 0.2-μm intervals with a QImaging Rolera-MGi Plus EMCCD camera and MetaMorph acquisition software (MDS Analytical Technologies). Images were processed using Fiji, and two-dimensional maximum intensity projections were created from the three-dimensional data sets when required.

### Structured Illumination Microscopy

3D stacks were collected at 0.125 um intervals on the Nikon N-SIM Structured Illumination Super-resolution Microscope with a 100x/NA 1.49 objective, 100 EX V-R diffraction grating, and an Andor iXon3 DU-897E EMCCD. Images were recollected and reconstructed in the “2D-SIM” mode (no out of focus light removal; reconstruction used three diffraction grating angles each with three translations).

### Transmission Electron Microscopy

Cytoskeleton preps were prepared as above and applied to 400 mesh formvar/carbon coated glow-discharged grids. Negative staining was performed by applying 1% phosphotungstic acid, pH 5.4, and grids were dried by blotting without washes. For thin sections, *Giardia* trophozoites that were pelleted or attached to ACLAR discs (Electron Microscopy Sciences) were fixed for 10 minutes in 4% paraformaldehyde and secondarily fixed for 1 hour in 1% osmium tetroxide. Cells were washed three times with cold ddH2O to remove fixative, dehydrated through ascending concentrations of ethanol (30%, 50%), and incubated for 1 hour in 2% uranyl acetate in 50% ethanol. Dehydration continued through 70% and 3 x 95% ethanol and was completed with three changes in 100% ethanol for a minimum of 10 minutes each change. Cell were embedded in 1:1 epoxy resin:acetone overnight at room temperature. The next day the resin was removed and replaced with 100% resin twice for two hours each. The ACLAR discs were placed at the bottom of a flat bottom beam capsule with the cells facing up and the capsule was filled with fresh resin. The blocks were polymerized at 70C overnight. The blocks were trimmed, and thin sections were cut with a Leica UCT ultramicrotome (Leica Ultracut UCT, Leica, Vienna, Austria) and stained with 2% uranyl acetate in 70% ethanol and lead citrate before viewing in the Talos L120C electron microscope (FEI Company/ThermoScientific, Hillsboro, OR., U.S.A. made in Eindhoven, The Netherlands) at 100KV. Images were acquired using the fully integrated Ceta CMOS camera.

## Results

### Biochemical fractionation of isolated cytoskeletons into disc and flagella-enriched fractions for shotgun proteomic analysis

Using cryo-ET of detergent extracted cytoskeletons (disc and flagella), Brown et al. recently defined specific regional variations in the ventral disc architecture, such as the ventral groove, disc margin and overlap zone areas (Brown et al., 2016). We hypothesized that the protein density disparities observed in these regions may represent targeting of specific DAPs to these disc regions. To find additional proteins that comprise these structurally distinct regions of the disc, we developed a novel biochemical fractionation protocol to disrupt the cytoskeleton and obtain a disc–enriched fraction, and used proteomic analysis and GFP tagging to identify candidate proteins.

Detergent extraction of *Giardia* trophozoites with Triton X-100 removes membrane and cytosol from the entire microtubule cytoskeleton and yields intact ventral discs and associated axonemes that are stable in PHEM buffer for weeks (Crossley and Holberton, 1983a; Crossley and Holberton, 1983b; Crossley and Holberton, 1985; Holberton and Ward, 1981). In a prior proteomic analysis of detergent-extracted, isolated cytoskeletons, we identified nearly twenty new DAPs that specifically localize to regions of the ventral disc or lateral crest (Hagen et al., 2011). Here we extracted trophozoites with both 1% Triton X-100 and 1M KCl (Figure 2A, P1 fraction), which facilitated the removal of contaminating proteins from the nuclei and cytosol as observed by negative staining EM. The disc and flagellar cytoskeletons were retained following the high salt extraction (Figure 2B, C). Negative staining of the P1 fraction (Figure 2D) showed the loss of the nuclei, and retention of the disc, flagella, and the funis. Subsequent extractions with Tris-EDTA and HEPES buffers were used to destabilize the cytoskeleton and release the disc from the axonemes and basal bodies, resulting in pellet or supernatant fractions (S2, S3, and P3). Using a strain with fluorescently tagged delta-giardin (Figure 2C, D), we determined that these additional treatments produced a fraction enriched for ventral discs (S3 fraction). These treatments also result in a relaxation of the normally domed and closed ventral disc spiral (Figure 2E, F). Many structural components of flagella are also removed in fraction S3 (Figure 2E, F). SDS-PAGE of each fraction resolved proteins enriched in each pellet (P1, P3) and supernatant (S2, S3) fraction (Figure 2G).

### Proteomics and mass spectrometry of fractions enriched in ventral disc or flagellar proteins

The composition of each fraction (P1, S2, S3, and P3) was determined by mass spectrometry. Proteins with fewer than five hits were excluded, and the four fractions were compared (Figure 2H and Supplemental Table 1). One hundred and fifty-seven proteins were identified with at least five hits in any fraction (Table 1). Tubulins, giardins, median body protein and GASP-180 family proteins were among the most commonly identified proteins. Thirty-five proteins occurred in every fraction examined, including 15 new and five previously described DAPs (e.g., beta giardin (Baker et al., 1988), delta giardin (Jenkins et al., 2009), SALP-1 (Palm et al., 2005)). Ten proteins were unique to the P1 fraction. Twenty-four proteins were enriched in the S2 fraction, and nine were enriched in the P3 fraction as compared to other fractions.

Tagging of selected proteins from these fractions allowed us to verify the selective enrichment of disc from flagellar components of the cytoskeletal preparations. Forty-nine DAPs were present in the P1, S2, or S3 fractions. While no proteins were specifically enriched in the S3 fraction (Figure 2H and Supplemental Table 1), 25 proteins in the S3 fraction had disc localization and only four localized to the flagella. In contrast, no DAPs were observed in fraction P3; the GFP-tagged proteins in this fraction localized only to the median body or the axonemes. Only one flagellar-localizing protein (DAP16996) was present in all four fractions. Twenty-two structural flagellar proteins (e.g., basal bodies or axonemes) were present in the P1, S2, or S3 fractions, and 16 were present in just the P1 or S2 fractions. Four lateral crest-localizing proteins were only present in the P1 fraction, and one lateral crest protein was present in both the P1 and S2 fractions. Thirty-nine DAPs that were previously identified or localized in an ongoing GFP tagging project (Nosala et al., 2018) were below the limit of detection in the proteomic analyses of the fractions. These 39 DAPs were included in subsequent analyses of ventral disc composition (Figure 3 and Figure 4, Supplemental Table 2).

**Figure 3.**
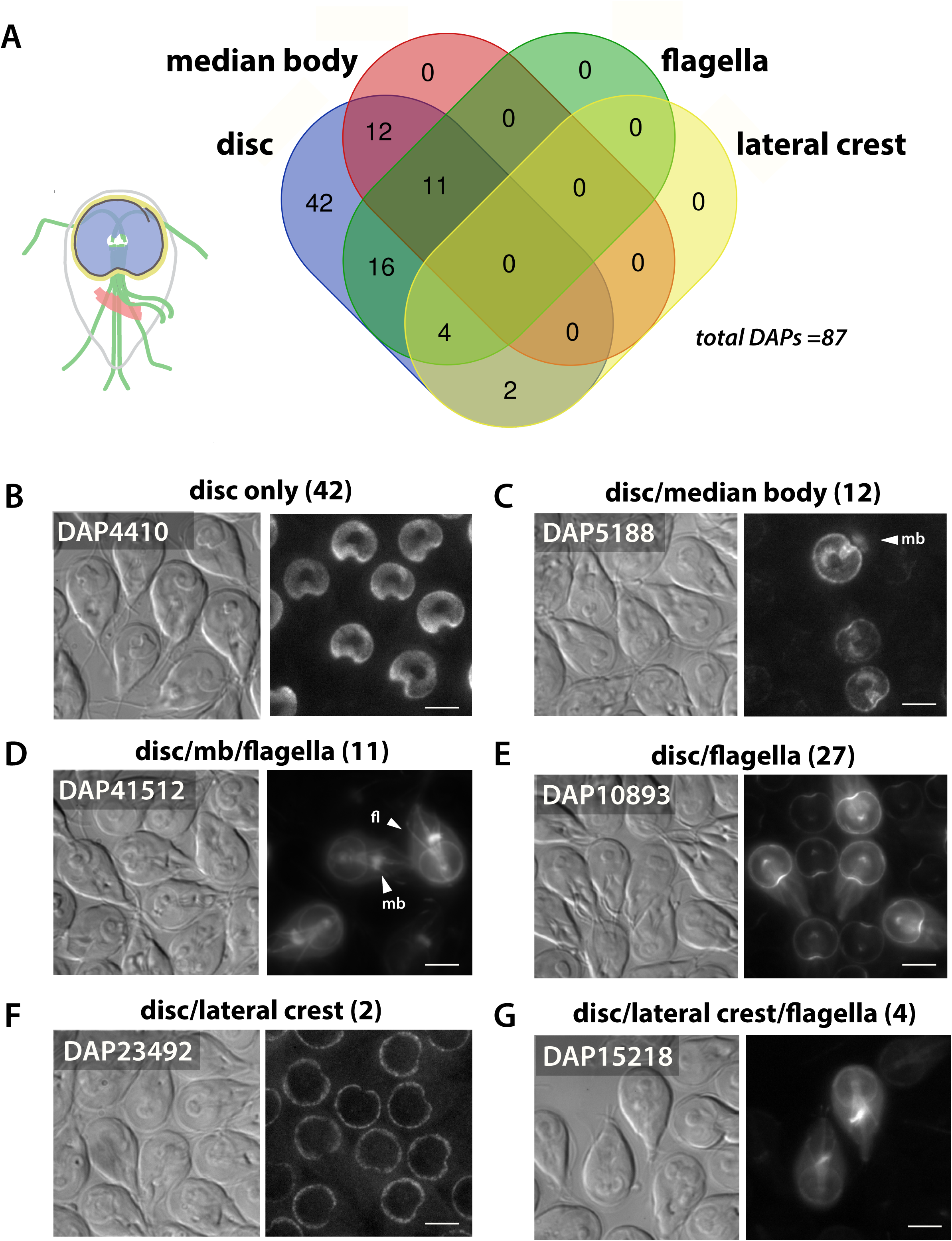
Many disc-associated proteins (DAPs) also localize to dynamic and stable microtubule structures. The subcellular GFP localizations of the 87 DAPs identified by fractionation and mass spectrometry (Figure 2) were categorized as localizing exclusively to the disc (N=42) (disc only, B) or to both the disc and other cytoskeletal structures, including the flagella (N=31), the median body (N=12), and lateral crest (N=4). The Venn diagram indicates DAPs with overlapping localizations to the disc, flagella, median body, and lateral crest (A). Representative localizations are shown for the 12 DAPs localizing to the disc and median body (mb, C); the 11 localizing to the disc, median body and flagella (mb and fl, D); and the 18 localizing to the disc and flagella only (N=18, E). Two DAPs localized to the lateral crest and disc (F) and two DAPs localized to the lateral crest and flagella (2). The flagella category includes localizations to either the basal bodies and/or to the cytoplasmic or membrane bound regions of the flagella. Scale bars = 5 μm.

**Figure 4.**
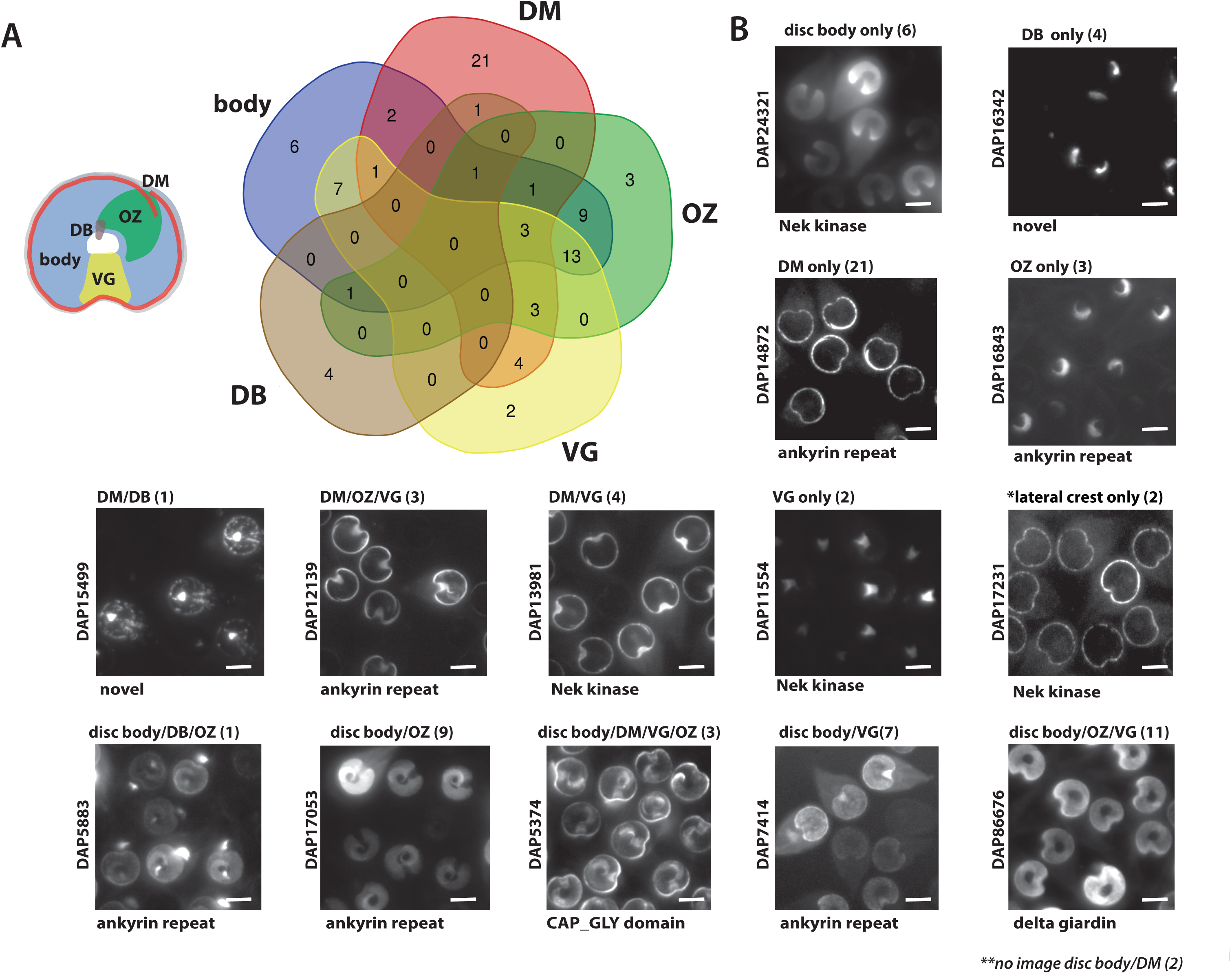
Disc-associated proteins localize to structurally distinct regions of the ventral disc. A Venn Diagram (A) shows categorization of the 54 new DAPs by their localization to distinct regions of the disc, including the MT spiral array (body), disc margin (DM), ventral groove (VG), dense bands (DB), and overlap zone (OZ). In B, representative DAPs defining each localization category are shown with their functional annotations. Scale bars = 5 μm.

### A revised inventory of disc-associated proteins

To determine the cellular localization of candidate DAPs identified here, we created C-terminal GFP tagged fusions of candidate DAPs and expressed them using their native promoters (Figure 3, Figure 4 and Supplemental Figures 1-4). The disc-specific localization of 54 new DAPs was confirmed, bringing the total number of verified DAPs to 87 (Figure 3 and Supplemental Table 2). Many of the newly identified DAPs localize to the disc and one other MT structure (Figure 3), such as the eight axonemes and basal bodies (27 total DAPs), the median body (23 total DAPs), or the lateral crest structure surrounding the disc (six total DAPs). Eleven DAPs localize to all of the primary MT-based structures including the disc, flagella, and median body. Forty-two of the 87 known DAPs localize only to specific regions of the ventral disc, and not to other cytoskeletal structures (Figure 3).

### DAP compositional variation in distinct regions of the ventral disc

The subcellular localizations of the 87 new and previously described disc- and median-body specific DAP-GFP strains were categorized into five distinct regions of the disc: the disc body (body), the disc margin (DM), the overlap zone (OZ), the ventral groove (VG), and the dense bands (DB) (Figure 4 and Supplemental Table 2). Overall, the majority of the 87 DAPs localize to more than one disc region, with 44 DAPs localizing to the disc body; 37 DAPs to the disc margin; 34 DAPs to the overlap zone; 33 DAPs to the ventral groove; and seven DAPs to the dense bands (Figure 4 and Supplemental Table 2). Thirty-six of the 87 DAPs localize to only one of the disc regions: six DAPS localize to the disc body only; three DAPs localize to the overlap zone only; 22 to the disc margin; four to the dense bands, and two DAPs only to the ventral groove. Two other DAPs localize only to the lateral crest and not to other disc or cytoskeletal elements (Figure 3, Figure 4 and Supplemental Table 2). There are also subtler yet noticeable variations in localizations within any particular disc region in the DAP-GFP strains; for example, some strains with disc body or disc margin localizations lack localization to the ventral groove region (Supplemental Figures 1-4).

### The majority of DAPs lack known MT-binding motifs

Of the total confirmed DAPs, only two possess known outer MT-binding motifs (Figure 5 and Supplemental Table 2). One is kinesin-6a (DAP102455), which localizes to the disc margin region and is the only kinesin associated with the disc. The other, DAP5374, has a CAP-GLY domain (Weisbrich et al., 2007) and localizes to four disc regions (Figure 5 and Supplemental Table 2). In contrast, 26 of the 87 DAPs lack homology to any known protein in other eukaryotes and localize to any of the five disc or lateral crest regions. An additional 29 DAPs localizing to any disc region simply contain ankyrin repeat domains (Figure 5 and Supplemental Table 2). Several DAPs are members of conserved protein families, including: three striated fiber (SF)–assemblins (beta-giardin, delta-giardin, and SALP-1 (Palm et al., 2003)); four annexins (e.g., alpha-giardins (Bauer et al., 1999; Peattie, 1990; Weiland et al., 2005; Weiland et al., 2003)), and 13 NEK kinases (Figure 5 and Table 1). Nine of the 13 NEKs lack conserved catalytic residues (Nosala et al., 2018). Other DAPs include those possessing a WD40 repeat domain (DAP15218), a DUF1126 domain (DAP41512), an Mlf1p domain (DAP16424), an ENTH domain (DAP3256) (Ebneter and Hehl, 2014), or SHIPPO repeat domains (DAP103164 and DAP9148).

**Figure 5.**
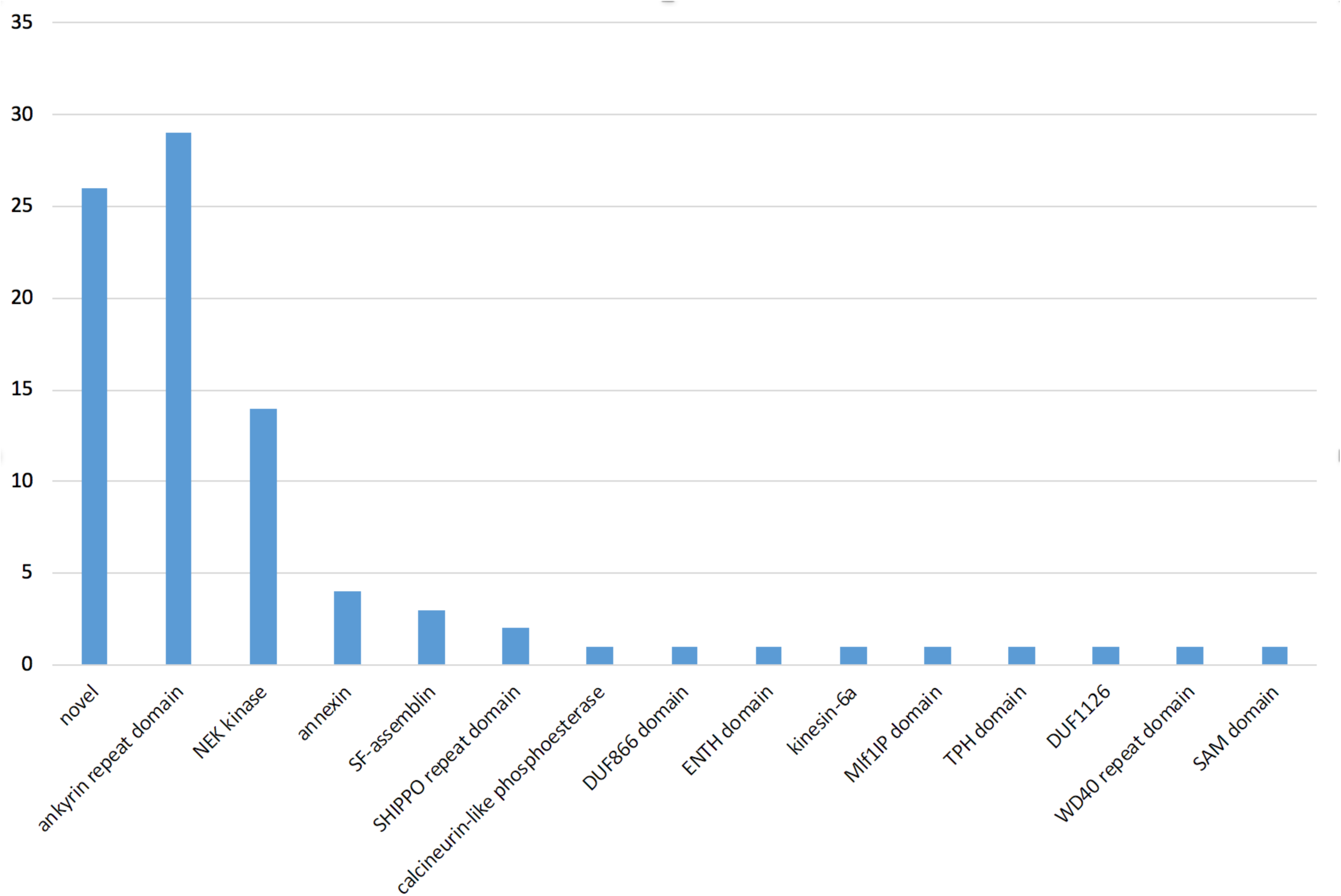
The majority of disc associated proteins (DAPs) lack known microtubule binding domains. PFAM domains are summarized and ranked by abundance for the 87 confirmed Giardia DAPs. CAP_GLY and kinesin motor domains are the only known MT-binding motifs. The five DAPs with Nek kinase domains that also possess ankyrin repeat domains are not included in the final summary.

### Ventral disc integrity is insensitive to microtubule drugs and high salt extraction

To evaluate the effect of MT stabilizing or depolymerizing drugs such as Taxol or nocodazole on disc structure, we treated cells with each of these drugs separately and quantified the area and overall structure of the disc body, overlap zone and central bare area. Treatment with either MT stabilizing or depolymerizing drugs had no effect on disc area or conformation (Figure 6A). In contrast, other cytoskeletal elements such as the flagella or median body are dynamic and sensitive to these MT drugs (Figure 6B).

**Figure 6.**
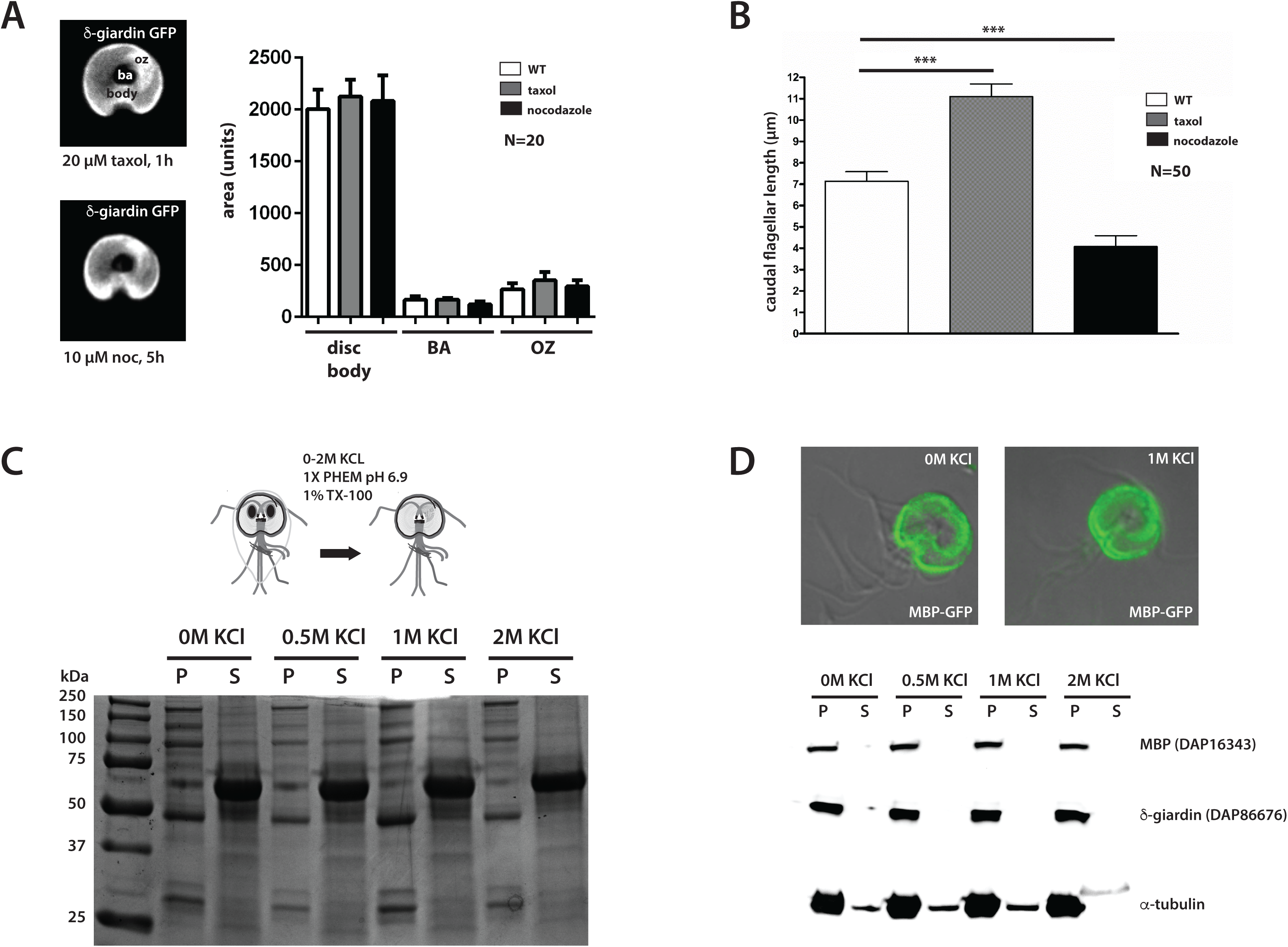
The overall composition and structural integrity of the ventral disc is insensitive to microtubule drugs and extraction with high salt and detergent. The area and overall structure of the disc body, overlap zone (oz) and central bare area (ba) are not affected by treatment with either MT stabilizing or depolymerizing drugs such as Taxol or nocodazole (A). In contrast, other cytoskeletal elements, including the flagella, are dynamic and sensitive to MT drugs (B). SDS-PAGE of fractions (pellet, P, and supernatant S) from 0 to 2M KCl extractions shows that salt also has little effect on extracted proteins (C). Furthermore, the GFP-tagged disc protein MBP remains associated with the disc even after extraction with 1M KCl (images in D, green), as do α-tubulin and the microribbon protein δ-giardin as seen via Western blot (D). Scale bars = 5 μm.

The detergent extraction of ventral disc cytoskeletons with up to 2M KCl had little effect on the recovery of disc-associated proteins as seen by SDS-PAGE (Figure 6C). Furthermore, the GFP-tagged disc protein DAP16343 (or MBP) remains associated with the disc even after extraction with 1M KCl. Alpha tubulin and delta-giardin, a component of the microribbons, also remain associated with the disc following high salt extraction (Figure 6D).

## Discussion

Cytoskeletal innovation and diversity are widespread in eukaryotic cells (Dawson and Paredez, 2013), and *Giardia* and other diverse emerging cell biological model systems offer a wealth of unexplored microtubule structures with unique functional properties (Russell et al., 2017). Here we identified and confirmed 54 new proteins in *Giardia’s* unique ventral disc, adding to the 33 previously described DAPs (Hagen et al., 2011). Eighty-five of the 87 DAPs lack homology to known MAPs that bind to the outside of MTs (Figure 5), and close to a third of DAPs lack any homology to proteins outside of *Giardia* species (Andersson et al., 2007). Through their binding to both the outside and inside of almost all disc MT protofilaments, many DAPs (including gMAPs and gMIPs) and protein complexes (e.g., MR-CB complexes) likely confer hyperstability to the disc singlet MT array (Brown et al., 2016; Schwartz et al., 2012). Other DAPs likely contribute to nucleating the spiral disc MTs, facilitating curvature and doming of the disc, or stabilizing MT plus and minus ends (Brown et al., 2016; Schwartz et al., 2012).

### Conserved MAPs and MIPs likely confer stability and flexibility to the disc singlet MTs

The singlet MTs of the curved disc spiral array create the domed shape of the disc that is crucial for *Giardia* attachment (Woessner and Dawson, 2012), and must withstand intense mechanical forces, much like the forces acting on ciliary doublet MTs during beating (Linck et al., 2014). Singlet MTs typically exhibit dynamic instability, although they can be stabilized or modulated by the activities of various plus-end binding proteins (e.g., EB1, XMAP215) (Akhmanova and Steinmetz, 2015; Bowne-Anderson et al., 2015). In contrast, the hyperstability of cilia, centrioles, and basal bodies is proposed to result from both the overall structure of the doublet and triplet MTs that comprise these organelles and from MAPs and MIPs associated along their entire lengths. Like hyperstable cilia, the ventral disc singlet MTs are coated with gMAPs, gMIPs and other associated protein complexes, forming a hyperstable spiral lattice that is resistant to detergent and high salt extraction (Figure 6).

More well-studied MAPs, like tektins or tau, stabilize MTs by binding to the outside surface of the microtubule (Amos, 2004; Amos, 2008). Giardia lacks both tektin and tau homologs (Morrison et al., 2007), and we identified only a handful of DAPs with known MT binding domains (Figure 5). However, several of the newly identified DAPs are homologous to proteins that may confer stability to axonemes in other ciliated cells (Supplemental Figure 2). Two DAPs (DAP9148 and DAP103164) are homologous to less well-studied SHIPPO repeat domain proteins predicted to stabilize and add rigidity to MTs of the sperm tail (Egydio de Carvalho et al., 2002). These two SHIPPO repeat DAPs localize not only to the disc, but also to the eight axonemes and the median body (Figure 3 and Supplemental Table 2), which implies a broader role for SHIPPO repeat proteins in MT stability. DAP16263 localizes to the disc overlap zone and cytoplasmic axonemes and belongs to conserved protein family (DIP13) that localizes to the centrioles and ciliary MTs in Chlamydomonas, possibly stabilizing MTs (Pfannenschmid et al., 2003). The human neuronal DIP13 homolog SSNA1 has recently been shown to be involved in MT nucleation and MT branching (Basnet et al., 2018). Thus, DAP16263 may contribute to stabilization, branching or assembly of disc MTs in the overlap zone region.

Single-particle cryo-ET of isolated doublet MTs identified microtubule inner proteins (or MIPs) that form a lateral and longitudinal meshwork in metazoan, algal, or Tetrahymena ciliary MTs (Ichikawa and Bui, 2018). Microtubule-stabilizing drugs like Taxol also bind to the inside of the microtubule wall (Nogales et al., 1999). This inner scaffold formed by MIPs is thought to strengthen tubulin dimer and protofilament coherence, promoting microtubule stability and/or elasticity during ciliary beating (Ichikawa and Bui, 2018). MIPs are also implicated in the stabilization and the elasticity of the singlet subpellicular microtubules of the malarial parasite Plasmodium, enabling the highly elastic, yet stable cytoskeleton of sporozoites to flex as parasites squeeze through host tissues (Cyrklaff et al., 2007).

As seen by cryoET, three microtubule inner protein densities (gMIPs 5, 7 and 8) are present in the lumen of the Giardia ventral disc singlet MTs, and are suggested to support the structural integrity and hyperstability of the disc (Schwartz et al., 2012). Although the molecular identities of periodically repeating gMIPs remain unknown, we confirmed that DAP41512 – a Rib72 homolog with a DUF1126 domain – localizes to the disc margin, as well as to the flagella and median body. Rib72 homologs are thought to be globular MIPs, and Tetrahymena RIB72A and RIB72B are essential for the assembly of A-tubule MIPs in cilia (Stoddard et al., 2018). FAP52 is another widely conserved MIP recently shown to localize to the inner junction of doublet MTs (Owa, 2018). Giardia has two FAP52 homologs (DAP15218 and DAP15996) that contain WD40 motifs and localize to all flagella. The localizations of Giardia Rib72 and FAP52 MIP homologs to the stable singlet MTs of the disc and the more dynamic median body MTs support a more general role for MIPs in promoting the stability and elasticity of interphase MT polymers that goes beyond stabilizing doublet and triplet MTs in axonemes.

### Novel and MR-CB complex DAPs may also create or stabilize the spiral, domed MT array

Some DAPs or disc protein complexes that likely regulate disc stability and architecture may be evolutionary innovations in Giardia. The MR-CB complexes in particular have been implicated in the stabilization of the domed disc conformation essential for proper parasite attachment (Nosala et al., 2018). Three microribbon DAPs are SF-assemblins, which associate with flagellar root structures in other protists (Weber et al., 1993), including the Toxoplasma apical complex (Francia et al., 2012). The identities of crossbridge or other MR DAPs remain unknown. Detergent-extracted discs are flattened, yet disc microtubules remained curved despite the loss of overall disc cohesion and crossbridge connections (Crossley and Holberton, 1983b) (Holberton and Ward, 1981). In our modified high salt and detergent extraction, we also observed the dissociation of crossbridge connections between the microribbons; furthermore, we confirmed that these hyperstable discs were flattened, and MTs retained their overall curvature (S3 fraction, Figure 2E, F). Thus, while intact MR-CB complexes are required for the overall domed architecture of the disc, the stable curvature of MTs in the spiral is likely governed by other DAPs (Holberton, 1981; Holberton and Ward, 1981).

Other abundant Giardia DAPs have less obvious roles in disc structure and stability. Thirty DAPs have ankyrin repeat domains, and five DAPs have both NEK domains and ankyrin repeat domains (Figure 4, Figure 5 and Supplemental Table 2). Ankyrin repeat proteins generally mediate protein-protein interactions and protein stability (Li et al., 2006). They are key components of the conoid MT organelle in the apicomplexan protist Toxoplasma gondii (Long et al., 2017). In human erythrocytes, muscles, and neurons, ankyrin proteins stabilize subsets of microtubules and can directly interact with tubulin in vitro (Bennett and Davis, 1981; Davis and Bennett, 1984).

### DAPs delineate discrete functional regions of the disc architecture

The disc MTs vary in length from 2 to 18 µm, and overall form more than 1.2 mm of polymerized tubulin (Brown et al., 2016). Almost all protofilaments of each disc MT are coated with different DAPs or complexes as they extend from the dense band nucleation zone and terminate at the disc margin (Brown et al., 2016). The strikingly distinct localization patterns of the 87 DAPs (Figure 4) mirror the structural regions in the disc defined by cryo-ET, including the disc margin, dense bands, ventral groove region and overlap zone (Brown et al., 2016).

Drugs that affect MT dynamic instability have little or no effect on disc MT dynamics or the overall disc structure ((Dawson et al., 2007) and Figure 6)) which implies that some DAPs may modulate dynamic instability at the plus or minus ends. In general, microtubule plus-end-tracking proteins (+TIPs) accumulate at growing plus ends, where they modulate and couple dynamic MT movements to cellular structures (Akhmanova and Hoogenraad, 2005). The plus ends of the ventral disc MTs are primarily located at the outer edge of the disc in the disc margin and at the edges of the overlap zone region (Brown et al., 2016). While more conserved plus end binding proteins like EB1 do not localize to the disc margin (Dawson et al., 2007), a large fraction of known disc proteins—thirty-eight DAPs—localize to the disc margin and other structures, with eight DAPs localizing exclusively to this region. The majority of disc margin DAPs have ankyrin repeat domains (12 DAPs), or NEK domains (six DAPs), and ten disc margin DAPs lack homology to any known protein. Thirteen disc margin proteins also localize to other MT-based structures including the flagellar axonemes, basal bodies (Lauwaet et al., 2011), or the median body (Supplemental Table 2), which implies that some disc margin DAPs have more general MT structural or regulatory properties.

Overlap zone DAPs may connect and stabilize the upper and lower portions of the MT spiral, maintaining the curved disc conformation required for attachment (Nosala and Dawson, 2017). Thirty-four DAPs localize to the overlap zone region, and a majority of these also localize to the disc body and/or disc margin regions. Sixteen overlap zone DAPs localize to the disc body, including the four known MR proteins. Two overlap zone DAPs also localize to the flagella and only one protein, DAP16843, localizes exclusively to the overlap zone region (Figure 4 and Supplemental Table 2). Morpholino or CRISPRi-based knockdowns of the overlap zone DAP16343 (or median body protein) have discs with flattened, open conformations (McInally SG et al., 2018; Woessner and Dawson, 2012), which suggests a role of the overlap zone region in establishing or maintaining the connections between the upper and lower regions of the disc (Figure 1).

The elasticity or flexibility of the disc MTs in regions such as the ventral groove or disc margin may result from differential binding of particular DAPs. In the ventral groove region, MTs lose much of their curvature and may be flexible (Brown et al., 2016). Two NEK kinases localize exclusively to this region. Four other ventral groove region proteins also localize to the disc margin, and the remainder localize to one or more other disc regions. Movements of the ventral groove and disc margin openings may mediate formation of the lateral crest seal, which would limit fluid flow underneath the disc in early stages of attachment (Nosala and Dawson, 2017). It remains unclear, however, how the disc margin and ventral groove DAPs mediate these movements and whether these movements modulate fluid flow (Nosala and Dawson, 2017).

Lastly, disc MT minus ends do not directly contact basal bodies but rather arise from a series of perpendicular bands termed the dense band (DB) nucleation zone (Brown et al., 2016). Three distinct dense bands in two regions consist of tightly packed microtubules that spiral into a nearly flat single plane. Although we have identified six DAPs localizing to these structures, the protein composition of the dense bands and the mechanism by which they support MT nucleation is undefined. About 39% of disc MTs nucleate from the disc margin (DM) region, possibly via a branching nucleation-type mechanism (Brown et al., 2016). One novel dense band protein (DAP15499) also localizes to the disc margin.

### Disc associated proteins may regulate dorsal daughter disc assembly and parental disassembly during division and encystation

During Giardia’s rapid mitosis and cytokinesis (Hardin et al., 2017), two daughter discs are assembled de novo on the anterior dorsal side of the parent cell, with the ventral sides of the new discs exposed on the cell surface (Tumova et al., 2007). During dorsal daughter disc assembly, the MT spiral is nucleated first, with the subsequent assembly and lengthening of the microribbons and crossbridges. The parental disc simultaneously undergoes disassembly and detachment.

Several putative regulatory proteins localize to the disc during division, including the sole Giardia aurora kinase (Davids et al., 2008), a protein phosphatase 2A (Lauwaet et al., 2007), and an ERK1 kinase that also localizes to the disc during encystation (Ellis et al., 2003). Two DAPs (DAP16424 and DAP5568) localize to both the disc margin and the basal bodies and may be associated with the Giardia polo-like kinase due to the presence of their respective Mlf1IP and DUF866 domains. Polo-like kinase is recruited to centromeres by phosphorylating a centromere scaffolding protein, PBIP1 (also called MLF1IP). Members of the NIMA-related kinase (or NEK) family regulate centrosome separation, spindle assembly and cytokinesis during cell division through targeted phosphorylation of proteins associated with the microtubule cytoskeleton (Fry et al., 2017). Giardia has an expanded repertoire of over 70 NEKs (Manning et al., 2011), and 12 NEKs localize to various regions of the disc (Supplemental Table 2). Two of the NEK kinases are putatively cell cycle-specific (Davids et al., 2011). Some of the disc-associated NEK kinases lack conserved catalytic residues, yet these putative NEK pseudokinases may still retain kinase activity (Reiterer et al., 2014). In addition to putative regulatory roles, the disc-associated NEKs may contribute to disc architecture and stability or even attachment dynamics.

### The molecular basis of functional diversity of Giardia’s singlet and doublet MT arrays

Different cellular contexts may dictate differential regulation of microtubule dynamics and stability. Like other protists, Giardia must modulate the dynamics and stability of various MT arrays in different regions of the cell during growth and cell division—the hyperstable ventral disc, the eight flagella, the median body, the funis, the caudal complex, and the two mitotic spindles all have differential stabilities and dynamics (Dawson, 2010). While Giardia’s eight axonemes have canonical doublet and triplet MTs, the disc, funis and caudal complex are stable interphase singlet MT arrays. In contrast, the median body and the tips of the flagella are dynamic and sensitive to MT drugs ((Dawson et al., 2007) and Figure 6).

How does Giardia simultaneously and differentially regulate these functionally diverse MT arrays? Conserved MAPs and MIPs may generally regulate MT stability or dynamics (Figure 3), whereas DAPs localizing to distinct regions of the disc likely confer more specific properties to the MTs in that region. One obvious way to distinguish ventral disc MTs and those of other arrays is by tubulin post-translational modifications (PTMs) (Garnham and Roll-Mecak, 2012). Tubulin PTMs may generate a functional diversity of MTs within Giardia by directing and/or recruiting specific DAPs or protein complexes to the specific MTs of the disc or axonemes. Giardia has a several tubulin tyrosine ligase-like (TTLL) homologs that may post-translationally modify tubulin (Morrison et al., 2007), and various types of tubulin PTMs (i.e., acetylation and glycylation) have been reported (Weber et al., 1997) (Campanati et al., 1999; Soltys and Gupta, 1994). Tubulin PTMs may also directly influence the stability of MT arrays or indirectly influence stability by regulating DAPs or other proteins that affect MT dynamics (Garnham and Roll-Mecak, 2012). We anticipate that the use of newly developed genetic tools such as CRISPR interference (CRISPRi) to repress both single or multiple endogenous genes in Giardia (McInally SG et al., 2018) will rapidly change how we investigate the roles of DAPs and PTMs in disc architecture, hyperstability, and attachment in this widespread parasite and emerging model system for cytoskeletal biology.

## Acknowledgements

This work was supported by NIH R01AI077571 to SCD and Center for Comparative Medicine at UC Davis School of Medicine T32 AI060555 Animal Models of Infectious Diseases Training Program to CN. We graciously thank Dr. Michael Paddy of the UC Davis MCB Microscopy Core for helpful advice and use of the SDC and SIM microscopes. We also thank Patricia Kysar of the UC Davis MCB Electron Microscopy Lab for advice and training on TEM and Michelle Salemi and Dr. Brett Phinney at the UCD Proteomics Core for help with mass spectroscopy.

